# Selective isolation of mouse glial nuclei optimized for reliable downstream omics analyses

**DOI:** 10.1101/2021.09.14.460386

**Authors:** Miguel A. Pena-Ortiz, Sarfraz Shafiq, Megan E. Rowland, Nathalie G. Bérubé

**Author notes:** Corresponding author: Nathalie G. Bérubé. These authors contributed equally to this work (Miguel A. Pena-Ortiz, Sarfraz Shafiq).

## Abstract

**Background:** Isolation of cell types of interest from the brain for molecular applications presents several challenges, including cellular damage during tissue dissociation or enrichment procedures, and low cell number in the tissue in some cases. Techniques have been developed to enrich distinct cell populations using immunopanning or fluorescence activated cell/nuclei sorting. However, these techniques often involve fixation, immunolabeling and DNA staining steps, which could potentially influence downstream omics applications.

**New Method:** Taking advantage of readily available genetically modified mice with fluorescent-tagged nuclei, we describe a technique for the purification of cell-type specific brain nuclei, optimized to decrease sample preparation time and to limit potential artefacts for downstream omics applications. We demonstrate the applicability of this approach for the purification of glial cell nuclei and show that the resulting cell-type specific nuclei obtained can be used effectively for omics applications, including ATAC-seq and RNA-seq.

**Results:** We demonstrate excellent enrichment of fluorescently-tagged glial nuclei, yielding high quality RNA and chromatin. We identify several critical steps during nuclei isolation that help limit nuclei rupture and clumping, including quick homogenization, dilution before filtration and loosening of the pellet before resuspension, thus improving yield. Sorting of fluorescent nuclei can be achieved without fixation, antibody labelling, or DAPI staining, reducing potential artifactual results in RNA-seq and ATAC-seq analyses. We show that reproducible glial cell type-specific profiles can be obtained in transcriptomic and chromatin accessibility assays using this rapid protocol.

**Comparison with existing methods:** Our method allows for rapid enrichment of glial nuclei populations from the mouse brain with minimal processing steps, while still providing high quality RNA and chromatin required for reliable omics analyses.

**Conclusions:** We provide a reproducible method to obtain nucleic material from glial cells in the mouse brain with a quick and limited sample preparation.

**Highlights:**

- Fast and easy isolation and sorting of glial nuclei from the mouse brain
- Reproducible and versatile processing of enriched nuclei for omics applications

## 1. INTRODUCTION

Glial cells regulate essential processes of the central nervous system during development and in the adult brain like myelination, immune response, dendritogenesis, neuromodulation, and synapse formation, maturation and elimination (Matejuk and Ransohoff, 2020; Vainchtein and Molofsky, 2020). Of the different glial cells, astrocytes, oligodendrocytes and microglia are the most studied, due to their different roles in the nervous system physiology and disease (Allen and Lyons, 2018). Astrocytes directly form signalling networks with neurons (Durkee and Araque, 2019) and are therefore integral to nervous tissue function, including synapse formation and maintenance (Allen and Eroglu, 2017). Microglia are the resident immune cells of the brain but also participate in neuronal development, synaptogenesis and synapse maturation, through pruning and the release of molecules including nerve growth factor and tumor necrosis factor (Matejuk and Ransohoff, 2020; Reemst et al., 2016). Oligodendrocyte precursor cells (OPCs) not only differentiate into oligodendrocytes responsible for myelination throughout life but are also able to form synapses with neurons and may establish bidirectional communication (Sakry et al., 2014, 2011). Oligodendrocytes form the myelin sheath, regulating conduction across the axon, but can also regulate synaptic transmission through potassium clearance and the release of factors like brain derived neurotrophic factor (Xin and Chan, 2020).

The differentiation, maturation and function of glial cells are tightly regulated by changes in their chromatin and transcriptome (Hammond et al., 2019; Lattke et al., 2021; Marques et al., 2018). Epigenetic changes during development direct cell fate and, after maturation, regulate cellular function through regulation of the transcriptome (Bayraktar et al., 2015; Eggen et al., 2019; Tiane et al., 2019). There is also increasing evidence that continuous and plastic communication exists between different types of glia throughout life, highlighting the need to understand cell type-specific contributions and changes (Domingues et al., 2016; Nutma et al., 2020; Vainchtein and Molofsky, 2020). The exploration of the transcriptome of glial cells and the correlation to chromatin architecture is therefore a key area of investigation that will expand our understanding of the role of these cells in physiological conditions and disease.

*In vivo* studies in the mouse brain provide several advantages over culture, the most salient being the ability to amalgamate all signals and their consequences in a complete system. However, *in vivo* molecular studies have been hampered by the heterogeneity of brain tissue, preventing the assignment of particular alterations to specific cell types (Maze et al., 2014). The size of the cell type population is also relevant for glial cells, which tend be fewer compared to neurons, or expression and chromatin differences might be confounded by higher levels in other cell types. Therefore, a necessary step for omics studies of brain cells is to purify, or at least enrich, for a cell type of interest prior to molecular analyses. To this end, several methods have been developed to enrich for particular cell types of interest, include sorting of immunolabelled cells or nuclei of the desired cell type (Douvaras and Fossati, 2015; Holt and Olsen, 2016; John Lin et al., 2017).

The first step in many protocols involves an enzymatic digestion to dissociate brain tissue into cell suspensions. The digestion time and type of enzymes must be carefully optimized to minimize loss of antigen targets of antibodies used in subsequent steps (Barres, 2014). Although these methods allow the isolation of whole cells, loss of membrane integrity from sheer stress can induce stress signals and could alter chromatin structure and gene expression profiles (Mo et al., 2015). Once a cell suspension is obtained, different approaches can be used to enrich for a specific cell type. Immunopanning is commonly used for this purpose and involves sequential plating of cell suspensions on antibody-coated plates to deplete cell populations based on membrane markers (Foo et al., 2011). This approach is most suitable to maintain and study these cell populations in culture; however, transcriptional changes are known to occur *in vitro*, and might not reflect the state of the cell prior to cell collection.

Alternatively, cells can be either labeled with fluorophore-tagged antibodies and isolated using fluorescence-activated cell sorting (Cahoy et al., 2008; Guez-Barber et al., 2012), or labeled with magnetically tagged antibodies and isolated with magnetic-activated cell sorting (MACS, (Marek et al., 2008). Although these methods allow good enrichment of populations, changes in transcription and chromatin structure could occur due to the initial dissociation steps and prolonged manipulation. Both FACS and MACS require incubation periods with antibodies, and sorting might require long periods of time or multiple elution rounds in the case of MACS (Chongtham et al., 2021). Another limitation is that each glial cell type is itself heterogeneous, making it challenging to find antibodies that will select for all sub-populations. (Marques et al., 2016; Matias et al., 2019). In order to limit disruptions caused by stress response inherent to the cell dissociation step, it is possible to instead quickly dissociate all cell membranes to isolate nuclei (Binek et al., 2019; Chongtham et al., 2021; Mo et al., 2015). Similar to whole cells, cell type -specific nuclei can be labeled with antibodies and sorted. While early studies simply used NeuN to obtain neuronal and non-neuronal populations (Jiang et al., 2008), specific cell-type nuclear markers have been used for oligodendrocytes (Mendizabal et al., 2019) and microglia (Nott et al., 2019). Although changes in the chromatin and transcriptome caused by signals from outside the nuclei would be removed, serial incubations and manipulation may still affect yield by breaking the nuclei.

To avoid the use of antibodies, genetic approaches emerged to tag cells or nuclei of interest with fluorescent proteins. The commercially available mice B6;129-Gt(ROSA)26Sor^tm5(CAG-Sun1/sfGFP)Nat/J^ (Mo et al., 2015), referred as Sun1-GFP, have a Cre-sensitive green fluorescent protein (GFP) and SUN1 nuclear lamina fusion protein that can be used to fluorescently tag nuclei of interest using cell-type specific Cre lines of mice (Mo et al., 2015). Available protocols for isolation and fluorescence activated nuclei sorting (FANS) require prolonged processing and manipulation before enrichment. To obtain nuclei suspensions, these methods use time-consuming ultracentrifugation, as long as 2.5 h in gradients (Percoll or iodixanol) that might be expensive or not readily available (Jiang et al., 2008). In addition, several protocols stain nuclei with dyes like DAPI or propidium iodide to help remove doublets during sorting that could interfere with downstream applications (Chongtham et al., 2021; Mo et al., 2015).

Here, we present a time saving procedure for isolation of fluorescent nuclei that uses a readily available sucrose gradient, and does not require fixation, ultracentrifugation or other prolonged manipulations and incubations. We show that this method is suitable for glial cells, providing high yield of nuclei with limited clumping. Furthermore, high enrichment of fluorescent nuclei using FANS was achieved by removing nuclei doublets without the use of dyes or additional antibodies. Finally, we also show that the enriched nuclei can be used for omics analysis, as RNA sequencing and assay for transposase-accessible chromatin followed by sequencing (ATAC-seq) revealed profiles characteristic of astrocytes, OPCs/oligodendrocytes and microglia. Our simplified workflow limits complications during nuclei enrichment, prevents possible artefacts in downstream applications and provides high quality RNA and chromatin for omics analyses

## 2. MATERIALS AND METHODS

### 2.1 Animals

All procedures involving animals were conducted in accordance with the regulations of the Animals for Research Act of the province of Ontario, Canada and approved by the University of Western Ontario Animal Care and Use Committee (2017-048). Mice were exposed to 12-hour light/12-hour dark cycles and fed *ad libitum* with tap water and regular chow.

The Sun1-GFP mice were obtained from the Jackson Laboratories (B6;129-Gt(ROSA)26Sor^tm5(CAG-Sun1/sfGFP)Nat/J^, IMSR JAX:021039, MGI:5614796) (Mo et al. 2015). Homozygous Sun1-GFP female mice were mated with male mice heterozygous for the tamoxifen-inducible Cre recombinase gene under the control of promoters specific for each glial cell type used: B6 Glast-CreER (Tg(Slc1a3-cre/ERT)1Nat, IMSR JAX:012586, MGI:4430111) for astrocytes (Slezak et al., 2007), Sox10-CreER (Tg(Sox10-icre/ERT2)388Wdr, IMSR JAX:027651, MGI:5634390) for OPCs (McKenzie et al., 2014), and CX3CR11-CreER (B6.129P2(Cg)-Cx3cr1^tm2.1(cre/ERT2)Litt^/WganJ, ISMR JAX:021160, MGI:5617710) for microglia (Parkhurst et al., 2013).

### 2.2 Genotyping

Genomic DNA from ear punches was extracted by incubating samples with DirectPCR Lysis Reagent (Viagen 102-T) and Proteinase K at 55°C overnight. DNA was amplified using primers specific to different genetically engineered alleles (Supplementary Table 1) using FroggaMix (FroggaBio FBTAQM). PCR conditions started with 95°C for 3 min, followed by 34 cycles of 95°C for 10s, 57°C for 20s and 72°C for 1min, ending with 72°C for 5 minutes.

### 2.3 Tamoxifen administration

Tamoxifen (10 mg, Sigma T5648) was mixed with 100 μl 95% Ethanol, incubated at 60°C until dissolved and diluted with 900 μl corn oil (Sigma C8267). Tamoxifen was injected intraperitoneally to induce Cre recombinase expression in different glial cell types. For OPCs and oligodendrocytes, Sun1-GFP lactating mothers crossed with Sox10-CreER males were injected with 2 mg tamoxifen for three consecutive days; for astrocytes, post-natal day 10 Glast-CreER;Sun1-GFP male pups were injected with 1 mg tamoxifen for three consecutive days; for microglia, 6 week-old CX3CR11-CreER;Sun1-GFP male mice were injected with 2 mg for five consecutive days.

### 2.4 Tissue collection and nuclei isolation

Mouse brains were dissected in ice cold phosphate buffered saline (PBS) and samples were frozen on dry ice before being stored at -80°C. Tissue was collected from tamoxifen-treated 20-day old Sox10-Cre;Sun1-GFP males (forebrain), 1 month-old Glast-Cre;Sun1-GFP male mice (cortex) and from 2 month-old Cx3cr1-Cre;Sun1-GFP male mice (cortex and hippocampus). For nuclei isolation, the solutions were made fresh before each experiment using DEPC water (diethyl pyrocarbonate, Sigma D5758) and kept on ice. Frozen tissue was homogenized in 500μl of homogenization buffer [*HB*, 20 mM Tricine KOH, 25 mM MgCl2, 250 mM sucrose, 1 mM DTT, 0.15 mM spermine, 0.5 mM spermidine, 0.1% IGEPAL-630, 1x protease inhibitor (Millipore Sigma 11873580001), 1 μl/ml RNaseOUT™ RNase inhibitor (Thermo Fisher Scientific 10777019)] with a pestle homogenizer for approximately one minute until a fine suspension was visible. The sample was then diluted to 7.5 ml with HB, filtered through a 40 μm strainer (Falcon 08-771-1) and carefully pipetted over 7.5 ml of cushion buffer [*CB*, 0.5 mM MgCl2, 0.88 M sucrose, 0.5 mM DTT, 1x protease Inhibitor (Millipore Sigma 11873580001), 1 μl/ml RNaseOUT™ RNase inhibitor (Thermo Fisher Scientific 10777019)]. Samples were centrifuged at 2800 x g for 20 mins at 4°C to pellet nuclei. The supernatant was carefully removed, and the pelleted nuclei were incubated for 10 min in resuspension buffer [*RB*, 500 μl 4% FBS, 0.15 mM spermine, 0.5 mM spermidine, 1 ul/ml RNaseOUT™ RNase inhibitor (Thermo Fisher Scientific 10777019) in PBS] and mixed by gentle pipetting.

### 2.5 Fluorescence activated nuclei sorting

Samples were filtered through a 20 μm strainer (pluriStrainer 43-10020-60) before sorting on a Sony SH800 Cell Sorter with a 100 μm nozzle, and the following gains for the sensors: FSC:14, BSC: 35%, FL1(EGFP):38%. Gating is initially set to discard larger events. Next, GFP intensity was plotted against GFP fluorescence area to eliminate doublets and nuclei clusters by discarding events with higher area. From this final gating step, events with lower GFP intensity were discarded to increase purity of the collected nuclei. Sample volumes sometimes needed to be adjusted with RB in order to maintain a constant flow of events per second. Sorting of the sample was paused throughout the session as needed to resuspend the sample and avoid clumping of the nuclei. For RNA extraction, sorting was also paused to mix the sorted nuclei with the lysis buffer in the collection tubes (described below). To assess the enrichment of GFP+ nuclei in the final sorted nuclei population, 5000 to 10000 nuclei were collected in PBS and an aliquot of the suspension was mixed with 1μg/ml DAPI and placed on a glass slide. The proportion of GFP+ nuclei was quantified with an inverted microscope (DMI 6000b, Leica) and digital camera (ORCA-ER, Hamamatsu).

### 2.6 RNA purification and RNA-seq library preparation

Sorted Sun1-GFP^+^ nuclei were collected in 100 μl lysis buffer from the Single cell RNA purification kit (NorgenBiotek 51800) with 2% β-mercaptoethanol. Immediately after sorting, additional lysis buffer was added to reach 600 μl of total lysate volume, as well as 600 μl 70% ethanol. RNA purification was performed according to the manufacturer’s instructions, including an in-column DNase treatment. Samples were eluted twice from the column with 8-12 μl of elution buffer. RNA concentration was determined using Qubit fluorometer (Invitrogen) according to the manufacturer’s instructions using 1-2 μl of sample. RNA integrity was determined using an Agilent Bioanalyzer. Libraries from microglia samples were constructed using 35 ng of RNA with the VAHTS Universal V8 RNA-seq Library Prep Kit for Illumina (Vazyme NR605-01), while astrocyte and OPC/oligodendrocyte libraries were made with 90 ng of RNA using VAHTSTM Total RNA-seq (H/M/R) Library Prep Kit for Illumina (Vazyme NR603-01) following the manufacturer’s instructions.

### 2.7 Assay for transposase-accessible chromatin (ATAC)

Approximately 150,000 sorted Sun1-GFP^+^ nuclei were collected in 50 μl of RNase free PBS. To concentrate the nuclei, the suspension was centrifuged at 3,000 rpm for 5 minutes at 4°C. The supernatant was then carefully removed, and nuclei were resuspended in tagmentation buffer containing Tn5 transposase (Vazyme TD501) for 45 min. at 37°C. Column purification of the samples was quickly performed with QIAquick PCR Purification Kit (Qiagen 28104) to end the tagmentation reaction. Enrichment and purification of the libraries was done according to the TruePrep DNA kit instructions. Quantitative PCR was used to determine the number of enrichment cycles to reach one third saturation, and all the libraries were amplified with more than 12 PCR cycles. Clean-up of all sequencing libraries was performed with SPRIselect beads (Beckman Coulter). The size and purity of the libraries were assessed using an Agilent bioanalyzer before storing at -80°C.

### 2.8 Deep sequencing and data analysis

Pooled equimolar libraries were sequenced at Canada’s Michael Smith Genome Sciences Centre (BC Cancer Research, Vancouver, BC, Canada, https://www.bcgsc.ca) using the Illumina Hiseq (Illumina Inc., San Diego, CA). The libraries were sequenced as a 150 bp paired end run. 50 million paired-end reads were obtained for RNA-seq and 100 million paired-end reads for ATAC-seq. Raw reads were pre-processed using Cutadapt and mapped against Mus musculus GRCm39 with HISAT2 for RNA-seq and bowtie2 for ATAC-seq. SAMtools was used to sort and convert SAM files. Gene abundance for RNA-seq was calculated using StringTie. ATAC-seq peaks were visualized using the Integrative genomic viewer IGV (Broad Institute). Heatmaps of differential gene expression from RNA-seq data were generated using Heatmapper (Babicki et al., 2016) with single linkage hierarchical clustering and spearman rank correlation. Expression enrichment of cell type-specific transcription factors was calculated using the cell type-enriched lists and method described in Zhang et al., 2014. Briefly, by dividing the average for each cell type by the sum of the averages of the other cell types.

## 3 RESULTS AND DISCUSSION

We used available transgenic mice to obtain brain tissue with GFP tagged nuclei in astrocytes, microglia or OPC/oligodendrocytes. Optimized nuclei isolation was used to obtain nuclei suitable for FANS (Figure 1A). Sorted nuclei were then used for RNA or chromatin extraction, followed by the corresponding procedures for RNA-seq and ATAC-seq (Figure 1B).

**Figure 1.**
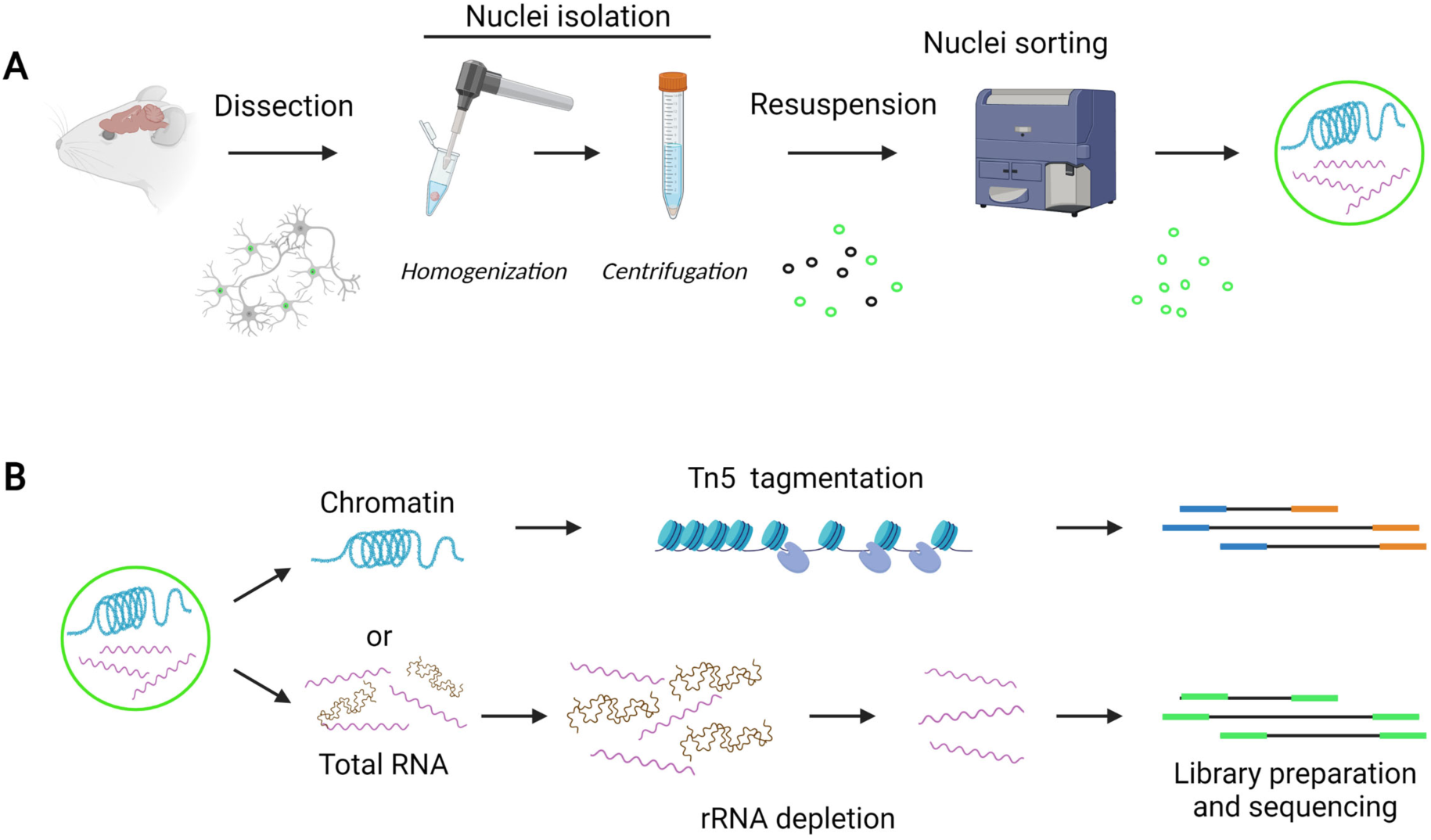
Protocol overview. (A) Tamoxifen-treated mice that express Sun1-GFP in astrocytes, OPC/oligodendrocytes or microglia are sacrificed, and brain tissue dissected and snap frozen. After tissue homogenization, nuclei are isolated by centrifugation and sorted to enrich for Sun1-GFP-tagged nuclei. (B) Sun1-GFP nuclei are immediately processed for total RNA extraction or chromatin accessibility assay (ATAC-seq) using Tn5. Ribosomal RNA is depleted from total RNA samples before generation of the RNA-seq libraries.

### 3.1 Nuclei isolation and sorting

We based this nuclei isolation protocol on the method previously published by (Mo et al., 2015). We observed loss of nuclei through homogenization and centrifugation steps, probably due to astringent conditions that affected nuclei integrity. To address this, we optimized the homogenization step and chose a nuclei pelleting centrifugation with sucrose gradient instead of a iodixanol gradient interlayer collection (Figure 2A). We modified the amount of detergent and length of dounce homogenization to maximize yield and integrity of the nuclei while limiting aggregation and clumping caused by leakage of the intranuclear material from ruptured nuclei (Maitra et al., 2021). To determine optimal detergent concentration and dounce time, frozen cortex tissue was homogenized in HB with 0.1%, 0.3% or 0.5% IGEPAL (Figure 2B). Samples were collected throughout the homogenization, stained with DAPI and quantified under a microscope. We observed that homogenization with 0.1% IGEPAL yielded the maximum number of nuclei with a dounce time of 105 s (Figure 2B and 2C). However, we observed that nuclei aggregation also increased with dounce time, potentially from rupture and leakage of a subset of nuclei. This clumping caused problems in the following steps, as it clogged filters during the filtration steps before sorting and ultimately decreased the final yield. Nuclei aggregates would also affect the sorting step, as they block the line or, if sorted, decrease purity. To prevent clumping of the nuclei, we used a dounce time of 1 min and pipetting was kept to a minimum with all nuclei suspensions. The nuclei yield obtained was enough for downstream applications with no clumping and minimal nuclei loss.

**Figure 2.**
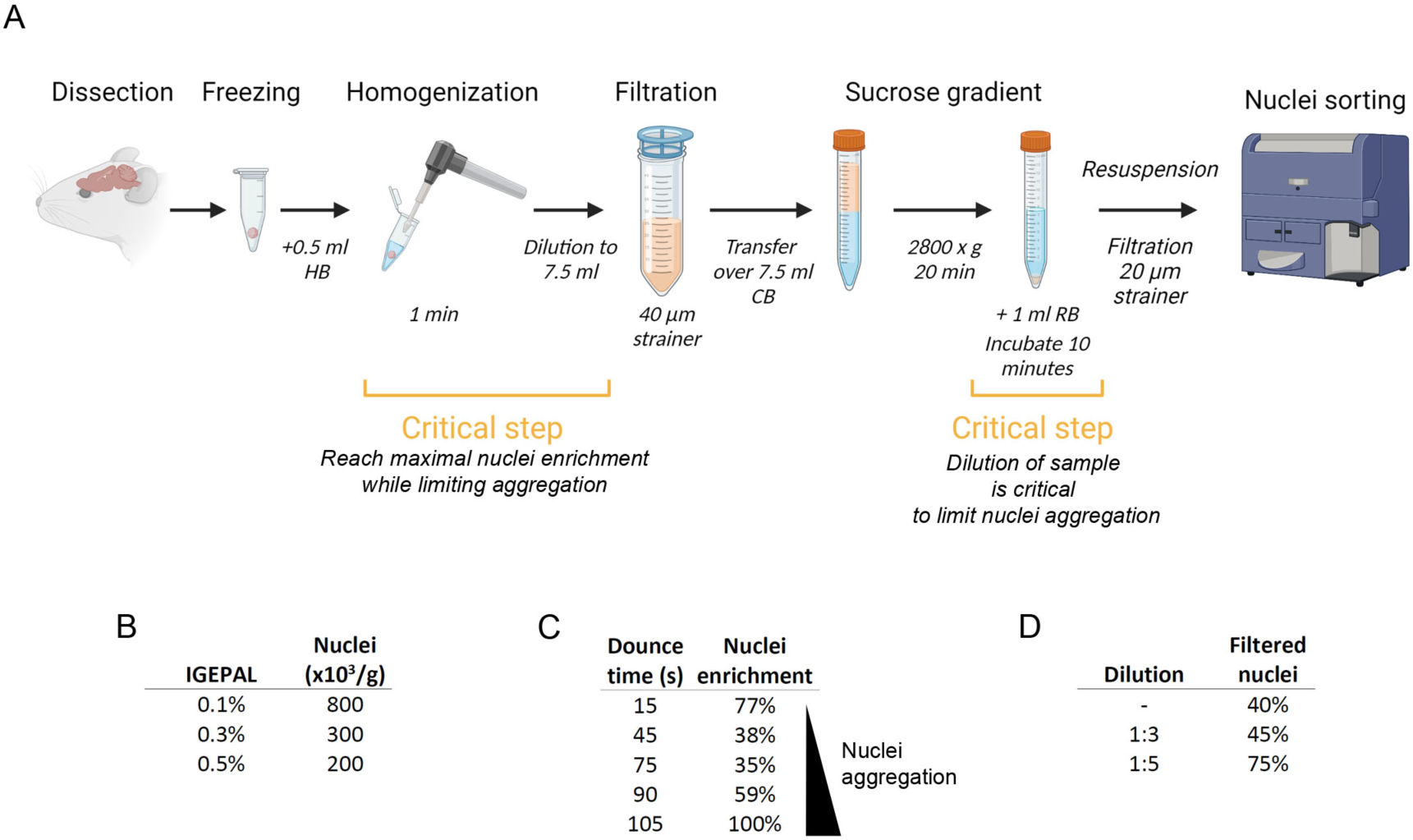
Nuclei isolation protocol. (A) Snap frozen brain tissue is homogenized with a pestle homogenizer. The sample is diluted before filtration and transferred to a sucrose gradient. Nuclei are pelleted through centrifugation, filtered and immediately sorted using a Sony SH800 Cell Sorter. (B) Maximum number of nuclei obtained throughout homogenization with different detergent concentrations. (C) Nuclei counts and aggregation increase with dounce time, GFP^+^ enrichment as percentage of the maximum number obtained. (D) Nuclei yield after filtration increases with dilution, enrichment as percentage of the nuclei counted before filtration.

In addition to detergent concentration and dounce time during homogenization, we found that a key step to obtain high yield was dilution of the nuclei suspension prior to filtration. The nuclei dilution at this step prevents clumping, clogging of the strainer and diminishes loss of nuclei. Furthermore, nuclei yield was significantly improved by increasing the dilution of the suspension to 1:5 (Figure 2D). We used an equivalent volume of a sucrose gradient (CB) and pelleted the nuclei, as we found this approach more efficient, cost-effective, and direct than using commercial gradients such as iodixanol. These gradients require ultracentrifugation and collection of the interlayer, which is difficult to visualize and obtain without leaving nuclei in the gradient, leading to reduced yield. Nuclei pelleting with a sucrose gradient circumvents these difficulties and allows nuclei resuspension in a buffer compatible with subsequent sorting steps. We also found that a 10 min incubation in buffer before resuspension limited nuclei rupture and clumping. Adding a filtration step just before sorting also decreased clumping issues (Figure 2A).

For FANS, we first gated the majority of events detected in the forward and back scatter channels, excluding larger events that could represent aggregated nuclei (Figure 3A). As an alternative to the use of dyes like DAPI to detect and remove doublets and smaller clusters, we plotted GFP signal intensity versus area using the population from the first gating (Figure 3B). Singlets appear in the plot in a linear distribution, which were selected to exclude events below the line (doublets) and close to the origin (GFP^-^). These settings identify the population of Sun1-GFP^+^ nuclei singlets in the suspension without any additional treatment or manipulation of the isolated nuclei. The gatings were optimized using GFP^-^ samples, to further determine the background levels of signal. Finally, to increase purity, a histogram of GFP signal was plotted, and only events with highest fluorescence signal were collected (Figure 3C).

**Figure 3.**
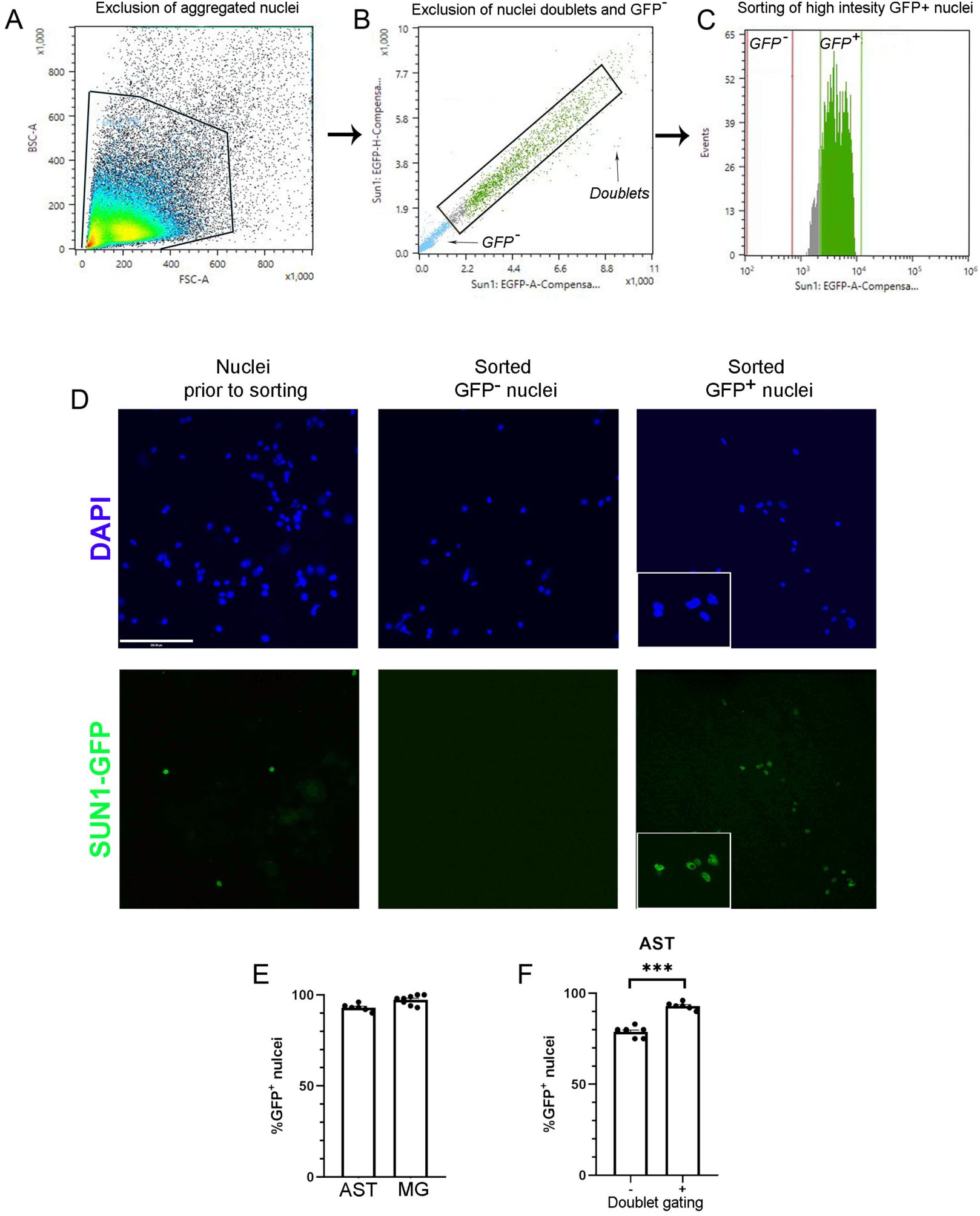
High enrichment of GFP tagged nuclei obtained through FANS. Representative images of the gating settings used to sort Sun1-GFP nuclei. (A) Gating set to discard clumps, events higher in the forward and back scattered channels; (B) population gated in A was then plotted according to EGFP signal intensity and area to identify nuclei singlets and remove doublets; (C) only nuclei singlets from the black box in B with higher EGFP signal intensity (shown in green) are collected, whereas events with lower EGFP signal (grey) are discarded. (D) DAPI-stained nuclei obtained before sorting or from sorted GFP^-^ and GFP^+^ collection tubes (E) Enrichment of Sun1-GFP^+^ sorted astrocyte (AST) or microglia (MG) nuclei (mean +/- SEM, n=6-8). (F) Enrichment of Sun1-GFP^+^ sorted nuclei as % of GFP tagged nuclei for astrocyte nuclei with or without gating to remove doublets, ***p<0.0001 unpaired t-test (n=6).

Samples collected before sorting and from GFP^+^ or GFP^-^ sorted nuclei were imaged to calculate the extent of Sun1-GFP^+^ nuclei enrichment. GFP signal was visible in Sun1-GFP sorted nuclei, and no GFP signal was found in the gating for GFP^-^ nuclei (Figure 3D). Our results indicate that this method allows enrichment above 90% for astrocytes and microglia nuclei (Figure 3E). We found that this high enrichment of GFP^+^ nuclei depends heavily on the absence of nuclei doublets or clusters. As an example, we determined enrichment for astrocyte nuclei with and without the second gating for doublets and found that the purity of Sun1-GFP nuclei improved by 10% when doublets and clusters were excluded (Figure 3F).

### 3.2 RNA extraction yield and quality

To maintain high RNA quality and yield, we paused sorting a handful of times to vortex the collection tube to ensure proper mixing of the nuclei in sheath solution with the lysis buffer. Collection in the lysis buffer facilitates downstream processing, as lysis of the nuclei starts as they are collected and does not require additional concentration steps before RNA extraction. We also added 2-mercaptoethanol (2% final concentration) to the collection tube to account for the volume of sheath solution containing the sorted nuclei. The RNA yield varied depending on the type of cell: 70 – 240 ng of RNA for astrocytes, 170 – 370 ng for OPCs/oligodendrocytes and 57 – 95 ng for microglia (Figure 4A). However, it is important to point out that these numbers cannot be compared directly because the samples used for each cell type were collected from different brain regions, at different time points. Moreover, each model uses distinctive Cre driver lines with presumably different Cre expression. The regimen of tamoxifen administration was also very different between animal models.

**Figure 4.**
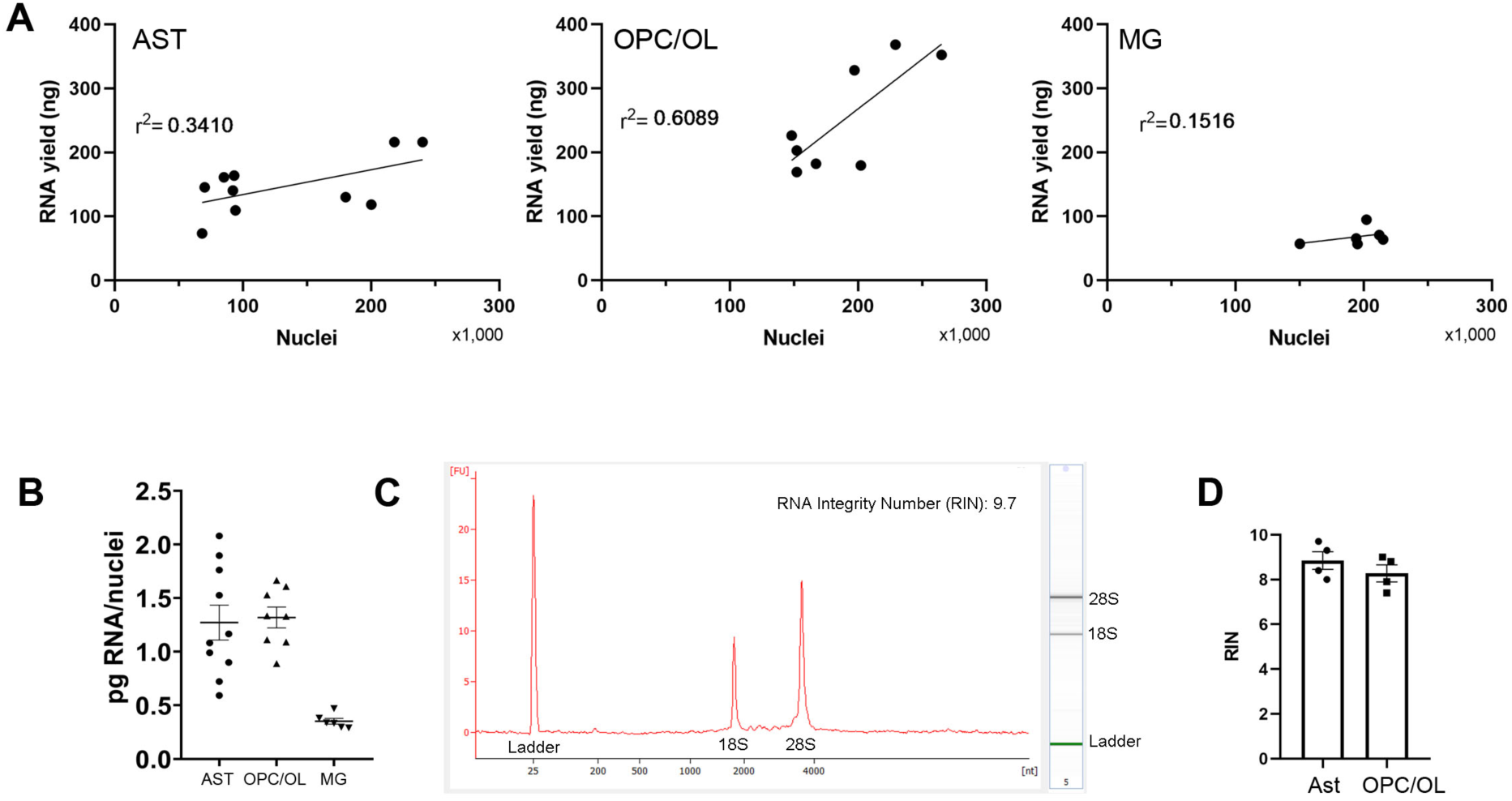
High RNA yield and quality from Sun1-GFP nuclei obtained through FANS. (A) Correlation between RNA extracted from the respective nuclei collected for astrocytes (AST) (p=0.0763), OPC/Oligodendrocytes (OPC/OLs) (p=0.0223) and microglia (MG) (p=0.4455). (B) Calculated RNA content per nuclei obtained for every sorting session for each cell type analyzed, ***p<0.0001 One-way ANOVA with Tukey’s Multiple comparisons (AST n=10, OPC/OL n=8, MG n=6). (C) Representative electrogram and gel view from Agilent Bioanalyzer used for RIN analysis. (D) Quality of RNA extraction as shown in RIN values of RNA samples extracted from astrocytes and OPC/OLs sorted nuclei (n=4).

Interestingly, in the case of astrocytes and microglia, the amount of RNA obtained appears to be determined by the cell type and not the number of nuclei. To better visualize the cell type-specific differences in the RNA yield obtained, we calculated the amount of RNA per nuclei and saw that microglia have significantly lower RNA content per nuclei and the results are less variable compared to astrocytes and OPC/Oligodendrocytes (Figure 4B). These results indicate that there might be cell type-specific differences in the RNA content in the nuclei of glial cells. To validate the quality of RNA extraction, we selected astrocytes and OPC/Oligodendrocytes for RIN analysis. Electrogram profiles and RIN from the Agilent bioanalyzer indicate that our protocol produced high integrity RNA (Figure 4C), as all RNA samples are above the quality threshold of a RIN above 7 for the glial cell types analyzed (Figure 4D). Therefore, despite differences in RNA content, enough high-quality RNA for further transcriptomic analysis can be obtained for each glial cell type with this method.

### 3.3 Cell type-specific gene expression and chromatin availability

We performed RNA-seq with three biological replicates using the RNA from sorted nuclei of astrocytes, microglia, and OPC/Oligodendrocytes. To further validate our approach, we compared the expression of well-known cell type-specific genes using the RNA-seq transcriptome data. Hierarchical clustering of these genes shows that indeed, the cell type-specific sets are enriched in their corresponding sample (Figure 5A). To further characterize our enriched populations, we used sets of transcription factors differentially expressed in each glial cell type, previously reported by Zhang et al., 2014 (Figure 5B), and observed distinct enrichment for the corresponding glial cell type and appropriate clustering of the samples. We also observe that astrocytes and OPC/Oligodendrocytes samples show higher similarity, compared to microglia. The distinct clustering of each cell type based on gene expression further validates that we obtained an enriched population of the desired glial cells, and that our optimized method provides a useful tool to analyze the transcriptome of enriched nuclei populations.

**Figure 5.**
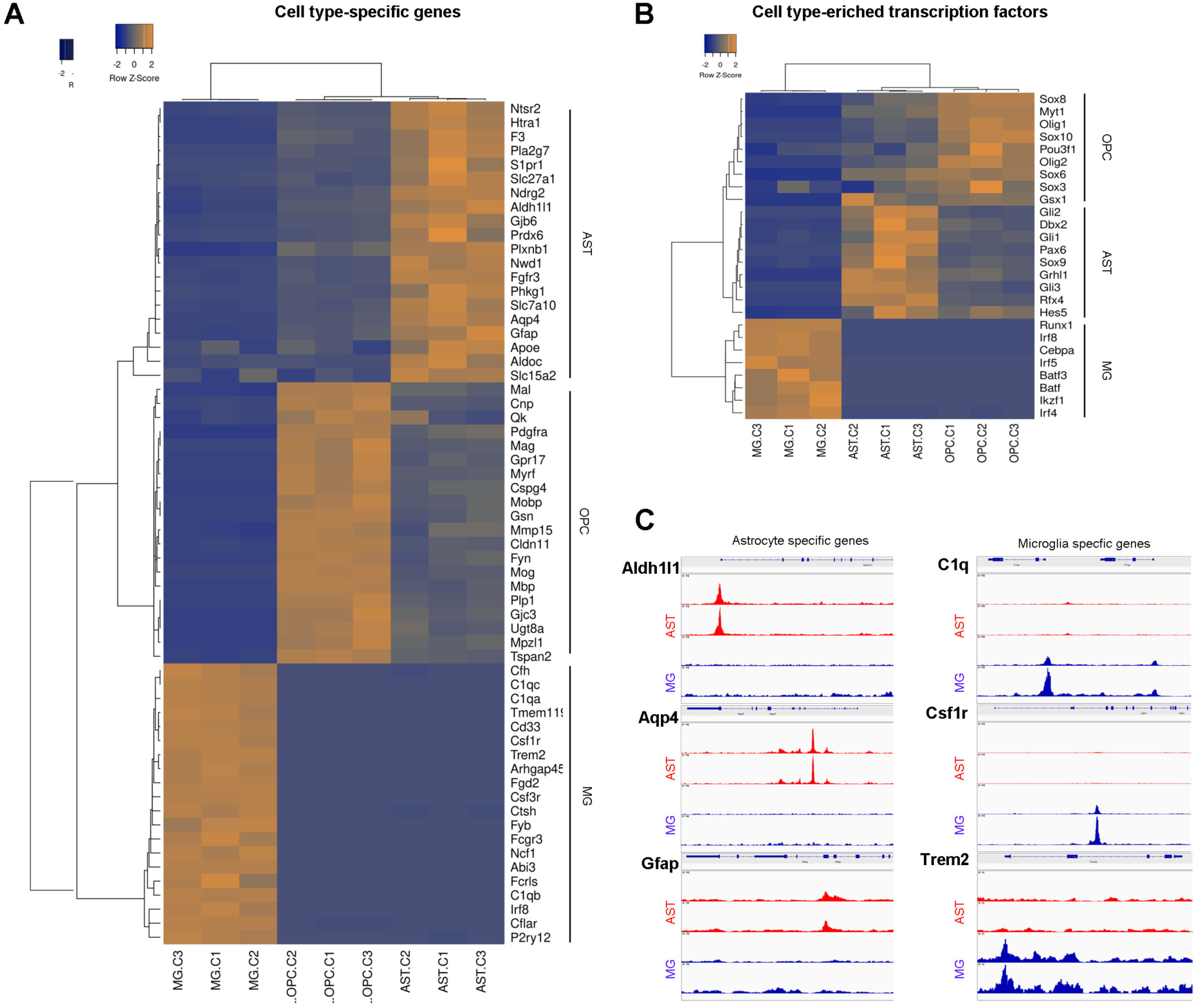
Differential expression and chromatin availability of cell type-specific genes determined by RNA-seq and ATAC-seq using sorted nuclei. (A) Heatmap of RNA-seq expression results with single linkage clustering and Spearman rank correlation shows higher expression of the corresponding cell type-specific genes for astrocytes (AST), OPC/Oligodendrocytes (OPC/OLs) and microglia (MG). (B) Enriched expression in the corresponding glial cell of transcription factors reported in (Zhang et al., 2014) to be cell type-enriched. (C) Chromatin availability from ATAC-seq analysis show distinct accessibility peak pattern in cell type-specific genes with differential expression determined with RNA-seq.

Astrocytes and microglia were selected for the validation of cell type-specific chromatin accessibility on sorted nuclei. We therefore performed ATAC-seq with two biological replicates using the sorted nuclei of astrocytes and microglia. To visualize the tracks of ATAC-seq, we selected cell-type specific genes that showed differential gene expression in our RNA-seq data. Consistent with cell type-specific gene expression, ATAC-seq also showed the distinct cell type-specific chromatin accessibility peaks for microglia and astrocytes in selected regions. For example, astrocytes specific genes *Aldh1l1, Aqp4* and *Gfap* showed more open chromatin in astrocytes but not in microglia. Similarly, microglia specific genes *C1q, Csf1r* and *Trem2* showed more open chromatin only in microglia (Figure 5C). Similar to the transcriptomic results, chromatin accessibility assays reveal patterns characteristic of enriched nuclei populations, confirming that our approach is suitable for omics analysis of the chromatin as well as transcriptome.

## 4. CONCLUSION

We present an optimized method for nuclei isolation and sorting of fluorescently tagged nuclei, applicable to a diverse population of cell types. Our method takes advantage of commercially available mouse lines that allow fluorescent labelling of nuclei from a variety of cell types of interest, avoiding the use of fixatives, antibodies or DNA dyes. The protocol used here for nuclei isolation and sorting consists of optimized, rapid and fewer manipulation steps, avoids cell dissociation stress while still yielding highly purified cell type-specific nuclei in sufficient quantity and quality for omics applications. This optimized protocol will therefore be a useful reference for the investigation of chromatin structure and gene expression in specific mouse brain cell types of interest.

## Abbreviations

AST: astrocyte
BSC: back scatter
DAPI: 4′,6-diamidino-2-phenylindole
DEPC: diethyl pyrocarbonate
DNA: deoxyribonucleic acid
DTT: dithiothreitol
FACS: fluorescence-activated cell sorting
FANS: fluorescence-activated nuclei sorting
FSC: forward Scatter
GFP: green fluorescent protein
MACS: magnetic-activated cell sorting
MG: microglia
OLs: oligodendrocytes
OPCs: oligodendrocyte precursor cells
PBS: phosphate buffered saline
RIN: RNA integrity number
RNA: ribonucleic acid

## Declaration of interest

None

## Funding

This study was supported by operating funds from the Canadian Institutes for Health Research to NGB (MOP142369). The funders had no role in study design, data collection and analysis, decision to publish, or preparation of the manuscript.

## ACKNOWLEDGEMENTS

We thank Dr. Frank W. Pfrieger for facilitating the use of the Glast-CreER mouse line and Alireza Ghahramani for analysis of the ATAC-seq data obtained from astrocyte and microglia nuclei.

---

**Supplementary table 1:**
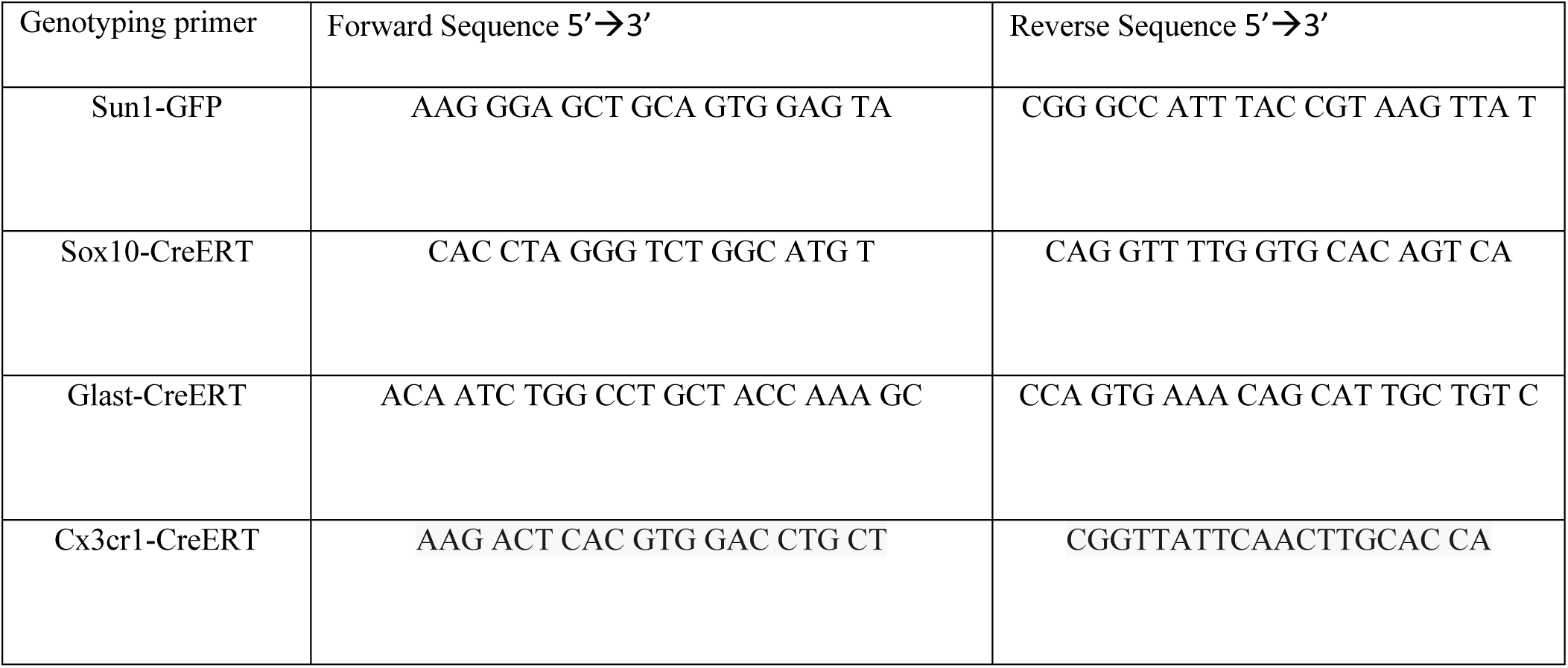
list of primer sequences.

## Notes

### Competing Interest Statement

The authors have declared no competing interest.

